# Sleep Loss Impacts the Interconnected Brain-Body-Mood Regulation of Cardiovascular Function in Humans

**DOI:** 10.1101/2021.04.27.441611

**Authors:** Adam J. Krause, Raphael Vallat, Eti Ben Simon, Matthew P. Walker

**Author notes:** Corresponding author Matthew P. Walker, **Email:**.

## Abstract

Poor sleep is associated with hypertension, a major risk factor for cardiovascular disease^1,2^. However, the mechanism(s) through which sleep loss impacts blood pressure remain largely unknown, including the inter-related brain and peripheral body systems that regulate vascular function^3^. In a repeated-measures experimental study of 66 healthy adult participants, we demonstrate four core findings addressing this question. First, a night of sleep loss significantly increased blood pressure—both systolic and diastolic, yet this change in vascular tone was independent of any increase in heart rate. Second, sleep loss compromised functional brain connectivity within regions that regulate vascular tone. Third, sleep-loss related changes in brain connectivity and vascular tone were significantly inter-dependent, with changes in brain nodes explaining the shift towards hypertension. Fourth, sleep-loss related changes in mood, specifically reductions in positive and amplification in negative states, each demonstrated an interaction with the impairments in brain connectivity and blood pressure. Together, these findings support an embodied framework in which sleep loss confers increased risk of cardiovascular disease through interactions between brain homeostatic control, mood-state and blood pressure.

## Main Text

### Introduction

Poor sleep is associated with hypertension, which is a primary risk factor for cardiovascular disease ^1,2^. Epidemiological studies indicate that both habitual short sleep duration and poor-quality sleep are associated with a greater incidence of hypertension, even when controlling for numerous other risk factors ^2,4^.

Prospective longitudinal studies have demonstrated that short sleep duration and worse quality sleep predict a higher future risk of hypertension ^2,5^. Manipulations using partial or total sleep deprivation causally and consistently increase blood pressure ^6,7^. Such evidence supports a possible link between the coinciding increase in sleep deficiency and increasing rates of cardiovascular disease in numerous first world nations^8^.

However, the mechanistic pathway(s) through which insufficient sleep increases blood pressure and thus cardiovascular disease risk remains unclear. This is especially true regarding a possible interaction between impaired central brain and peripheral body systems that are known to control vascular blood pressure. Indeed, the brain plays a causal role in the regulation of peripheral physiology, including that of blood pressure and heart rate ^3^. Core to this control are a discrete set of viscerosensory brain regions that that map and also instigate changes in vascular function.

Defining this cortical and subcortical network have been animal models demonstrating that direct stimulation of the amygdala ^9^, the cingulate ^10^, the insula ^11,12^ and medial prefrontal cortex (MPFC) ^10^ all elicit changes in blood pressure. In humans, activity within, and connectivity between, the amygdala, cingulate, and insular cortices ids in the regulation of arterial blood pressure, as well as event-related reactivity of vascular blood pressure ^13^.

Of this collection of regions, the insular cortex and MPFC are of particular importance in the regulation of vascular state. Both regions are involved in the central brain regulation of autonomic state, including the mapping of current autonomic balance and modulating autonomic outflow ^3,11,14^. Activity in the insula and MPFC have further been linked with increased vasoconstriction in subjects with a history of coronary artery disease ^15^, and ischemic injury to the insula induces endothelial dysfunction and inflammation ^16^. Similarly, lesioning of the MPFC increases plasma levels of adrenocorticotropic hormone and corticosterone ^17^, both important stress-response factors which can impact cardiovascular function ^18^. The insula and MPFC are unique in the regulation of cardiovascular function through direct connections to brainstem areas ^19,20^. Finally, the brain, including the insula and MPFC, can further impact vascular function through its influence on immune function as vascular endothelial cells are sensitive to inflammatory processes ^21,22^.

Although the peripheral autonomic nervous system is a known regulatory pathway for vascular and endothelial function control ^23^, how the autonomic nervous system interacts with these central brain viscerosensory networks under conditions of sleep loss, resulting in a shift towards hypertension, remains uninvestigated.

Building on this collective evidence, and addressing this unresolved question, here, we sought to test the hypothesis that one pathway through which insufficient sleep instigates a shift from normotensive towards a hypertensive increase in blood pressure is through an altered relationship with central functioning of brain vascular control networks and the peripheral body cardiovascular system. Specifically, we tested the prediction that elevations in blood pressure caused by a night of sleep deprivation are significantly associated with changes in the functional connectivity state of the above-described a priori vascular control networks of the human brain, indicative of a brain-body mechanistic inter-relationship.

In short (and see Methods for details), 66 healthy adult participants who were otherwise normotensive, were enrolled in a counterbalanced, repeated-measures design involving two conditions: one night of sleep (Sleep Rested) and one night of sleep loss (Sleep Deprivation). In each condition, participants had their cardiovascular state, including blood pressure and heart rate, assessed at circadian-matched morning timepoints, and received a resting-state functional magnetic resonance imaging (fMRI) scan following sleep or sleep deprivation—the latter being a recognized method for mapping the state of network connectivity of the human brain ^24^.

### Results

#### Sleep Loss & Cardiovascular Function

We first tested the hypothesis that sleep deprivation significantly alters key vascular, cardiac and autonomic metrics linked with cardiovascular disease^25^: systolic and diastolic blood pressure, heart rate, and heart rate variability.

Supportive of the hypothesis and prior findings^1,2,4,5^, sleep deprivation significantly increased systolic blood pressure, relative to the same individuals under sleep rested conditions (**Fig. 1A**; Systolic: sleep-rested mean = 107.6 mmHg ± 11.8 SD, sleep-deprived mean = 112.1 mmHg ± 12.8 SD, p = 0.003). This was similarly true of diastolic blood pressure, which also increased significantly following sleep loss (**Fig. 1A**; Diastolic: sleep-rested mean = 71.1 mmHg ± 7.3 SD, sleep-deprived mean = 73.7 mmHg ± 7.9 SD, p = 0.03).

**Figure 1.**
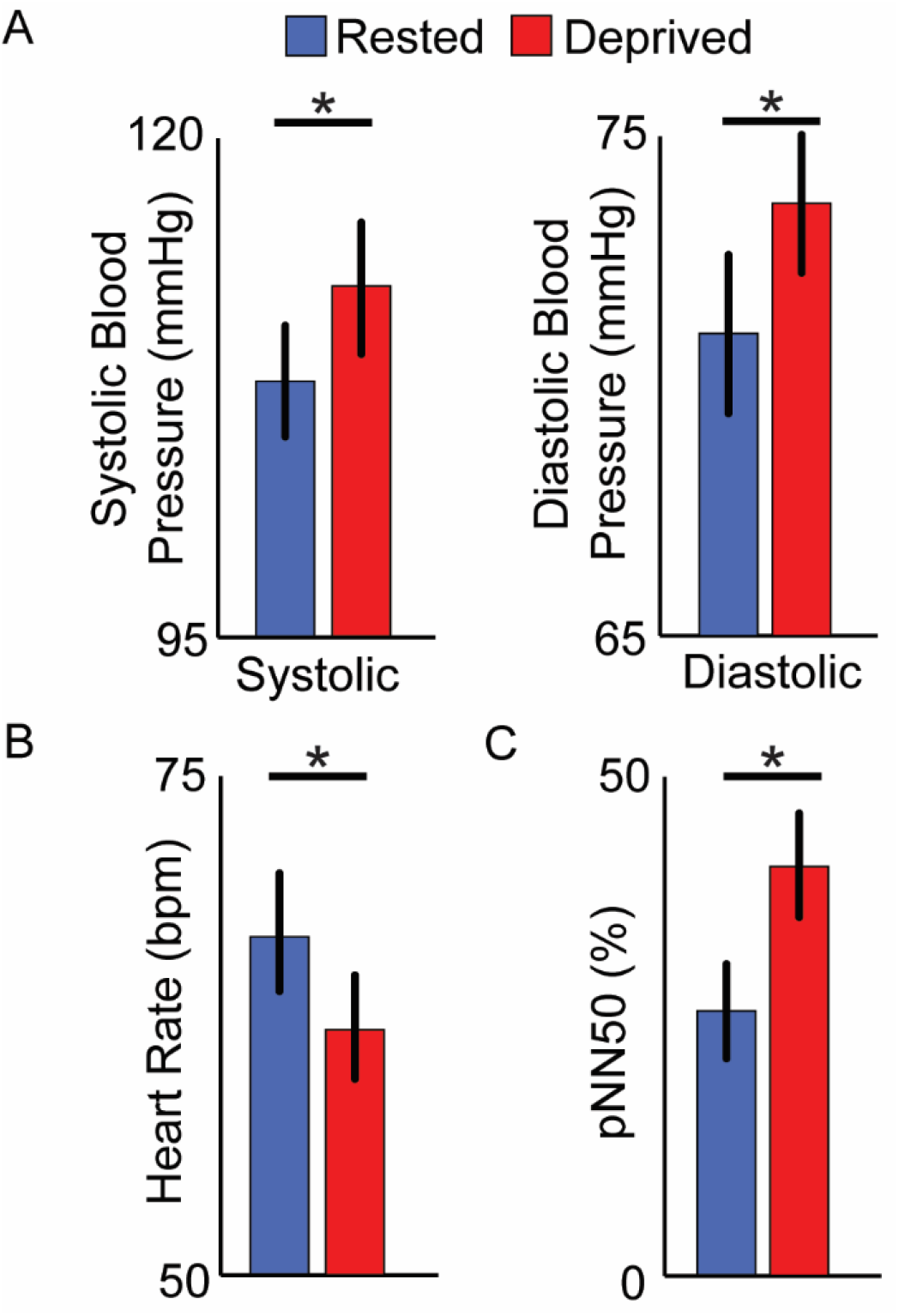
Cardiovascular outcomes. **A**, Following sleep deprivation, there was an increase in both systolic and diastolic blood pressure, relative to the sleep rested condition. **B**, Sleep deprivation resulted in a reduction in heart rate, relative to the sleep-rested condition. **C**, Sleep deprivation resulted in an increase in heart rate variability (pNN50), relative to the sleep-rested condition. Error bars indicate SEM. \**p < 0*.*05*.

These changes in blood pressure occurred in the absence of an increase in heart rate. Specifically, there was an overall reduction of resting heart rate under conditions of sleep loss, relative to the sleep rested condition (**Fig 1B**; sleep-rested mean = 67.5 bpm ± 11.6 SD, sleep-deprived mean = 62.9 bpm ± 10.2 SD, p < 0.001), a finding that is consistent with previous reports regarding sleep loss^6,26-29^, but see^30^.

Corresponding to the decrease in heart rate, there was an increase in heart rate variability under conditions of sleep deprivation, relative to the sleep-rested condition (**Fig 1C**; pNN50: sleep-rested mean = 27.5% ± 20.5% SD, sleep-deprived mean = 42.0% ± 19.1%, p < 0.0001). Since the pNN50 reflects vagal modulation of the heart^31^, this observed increase in heart rate variability under conditions of sleep loss is consistent with increased parasympathetic input to the heart linked to the state of extended wakefulness^26,28^.

Further advancing a dissociation between increased peripheral vascular blood pressure in the absence of accelerated heart rate, there was no significant association between the differential changes in blood pressure and heart rate between conditions ([sleep-rested] – [sleep deprivation]) for either systolic or diastolic measures (**Fig 2A,B**; Systolic: r = 0.06, p = 0.61; Diastolic: r = 0.03, p = 0.84). A similar lack of association was observed between the sleep-loss changes in heart rate variability and in blood pressure (**Fig 2C,D**; Systolic: r = −0.09, p = 0.48; Diastolic: r = −0.05, p = 0.69). In contrast, there was a significant association between the sleep-loss related reduction in heart rate and the corresponding increase in heart rate variability (r = −0.48, p < 0.0001).

**Figure 2.**
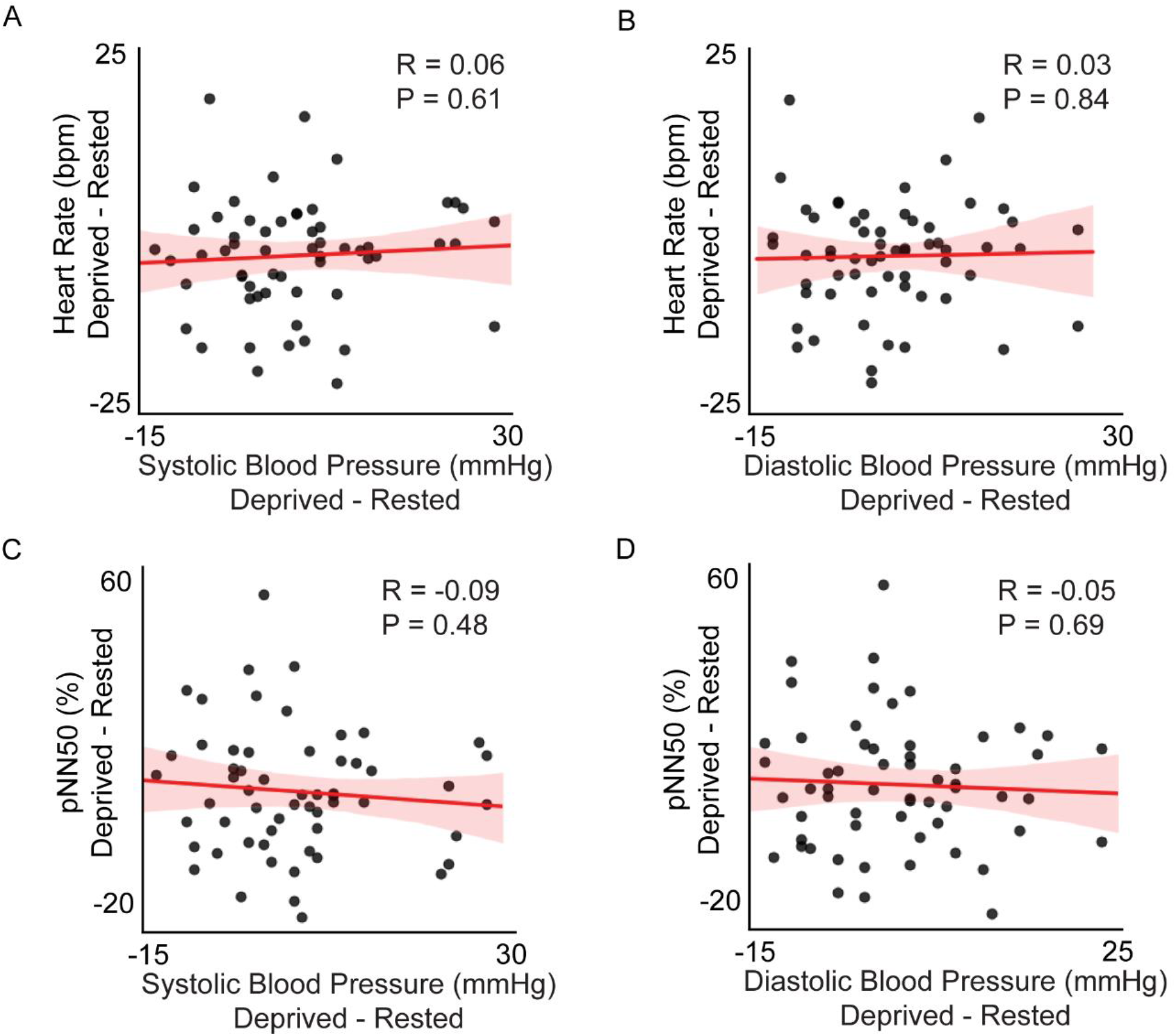
Associations between cardiovascular outcomes. Scatterplots represent non-significant correlations between sleep-loss changes (deprived – rested) in cardiovascular outcomes, including (**A**) heart rate and systolic blood pressure, (**B**) heart rate and diastolic blood pressure, (**C**) heart rate variability (pNN50) and systolic blood pressure, and (**D**) heart rate variability (pNN50) and diastolic blood pressure.

#### Sleep Loss & Viscerosensory Resting Brain Connectivity

Next, we assessed functional connectivity within the a priori network known to be associated with cardiovascular control in humans and non-human primates, including the amygdala, anterior cingulate, insula, and ventromedial and medial prefrontal cortices^3,9-13^. Within this network, twelve connections were found to be significantly altered following FDR-correction for multiple tests under conditions of sleep deprivation, relative to the sleep rested state (q = 0.05)^32^. These included connectivity pathways between the insula and anterior cingulate, the insula and MPFC, the anterior cingulate and the MPFC, the anterior cingulate and the amygdala, the ventromedial frontal and MPFC, and MPFC and amygdala (**Fig 3A,B, Table S1**).

**Figure 3.**
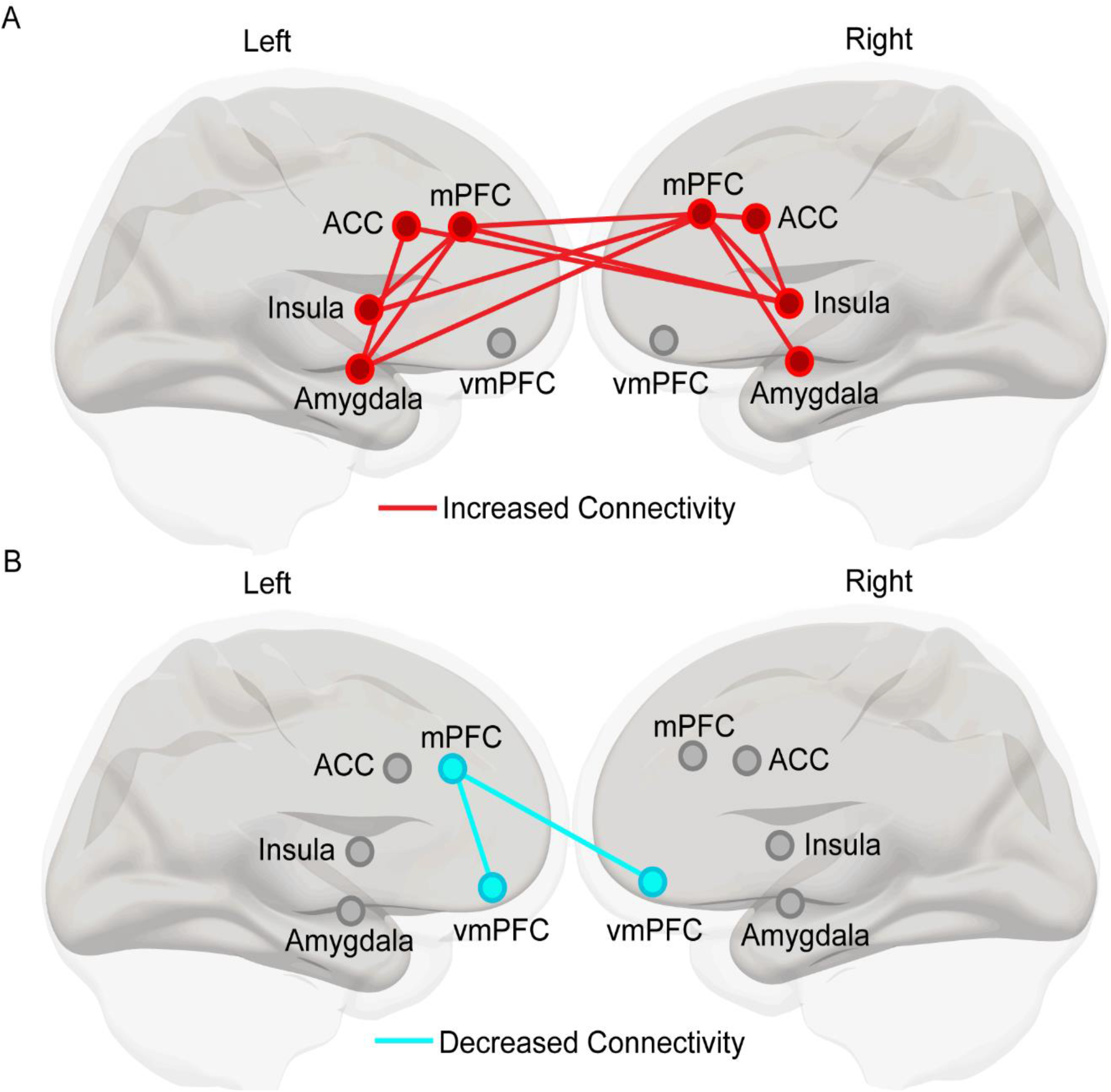
Changes in brain connectivity in regions of interest. Connections shown are significant following FDR-correction (q = 0.05). **A**, Connections that are significantly increased in the sleep deprived condition relative to the sleep-rested condition. **B**, Connections that are significantly decreased in the sleep-deprived condition relative to the sleep rested condition.

Of these twelve, there was a significant reduction in connectivity between the ventromedial frontal and MPFC following sleep deprivation, relative to a night of normal sleep (t(65) = 3.2, p = 0.008, FDR-corrected, **Fig. 3B**). The remaining eleven region of interest (ROI) pairs demonstrated a significant increase in their connectivity (**Fig. 3A, Table S1**), with the connectivity changes all jointly sharing at least one of two nodes: the insula and MPFC. First, the insula had significantly increased connectivity with the anterior cingulate cortex (p = 0.01, FDR-corrected). Second, there were increases in connectivity between the insula and the MPFC (p< 0.0008, FDR-corrected). The MPFC demonstrated increased connectivity to both the anterior cingulate in the right hemisphere (p < 0.0001, FDR-corrected), and amygdala in both hemispheres (p < 0.02, FDR-corrected). Details of these connectivity changes and their significance are described in **Supplement**.

#### Sleep Loss, Cardiovascular Function & Viscerosensory Resting Brain Connectivity

We next tested whether these two sets of identified changes—alterations in brain connectivity and alterations in peripheral blood pressure were significantly inter-related. First, greater sleep-loss-related increases in insula connectivity were significantly associated with a greater rise in systolic blood pressure, especially between the insula and the MPFC (r(60) = 0.25; p = 0.05; **Fig. 4A**). Second, sleep-loss-related decreases in MPFC connectivity with the amygdala was significantly related to increases in diastolic blood pressure (r(61) = −0.29, p = 0.02; **Fig. 4B**). Therefore, sleep-loss changes in systolic and diastolic blood pressure were associated with alterations in brain connectivity, with a clustering around changes in insular and MPFC connectivity, respectively. Unlike the association between brain and blood pressure, no associations between sleep-loss related changes in heart rate or heart rate variability and changes in functional connectivity were observed (all p > 0.05).

**Figure 4.**
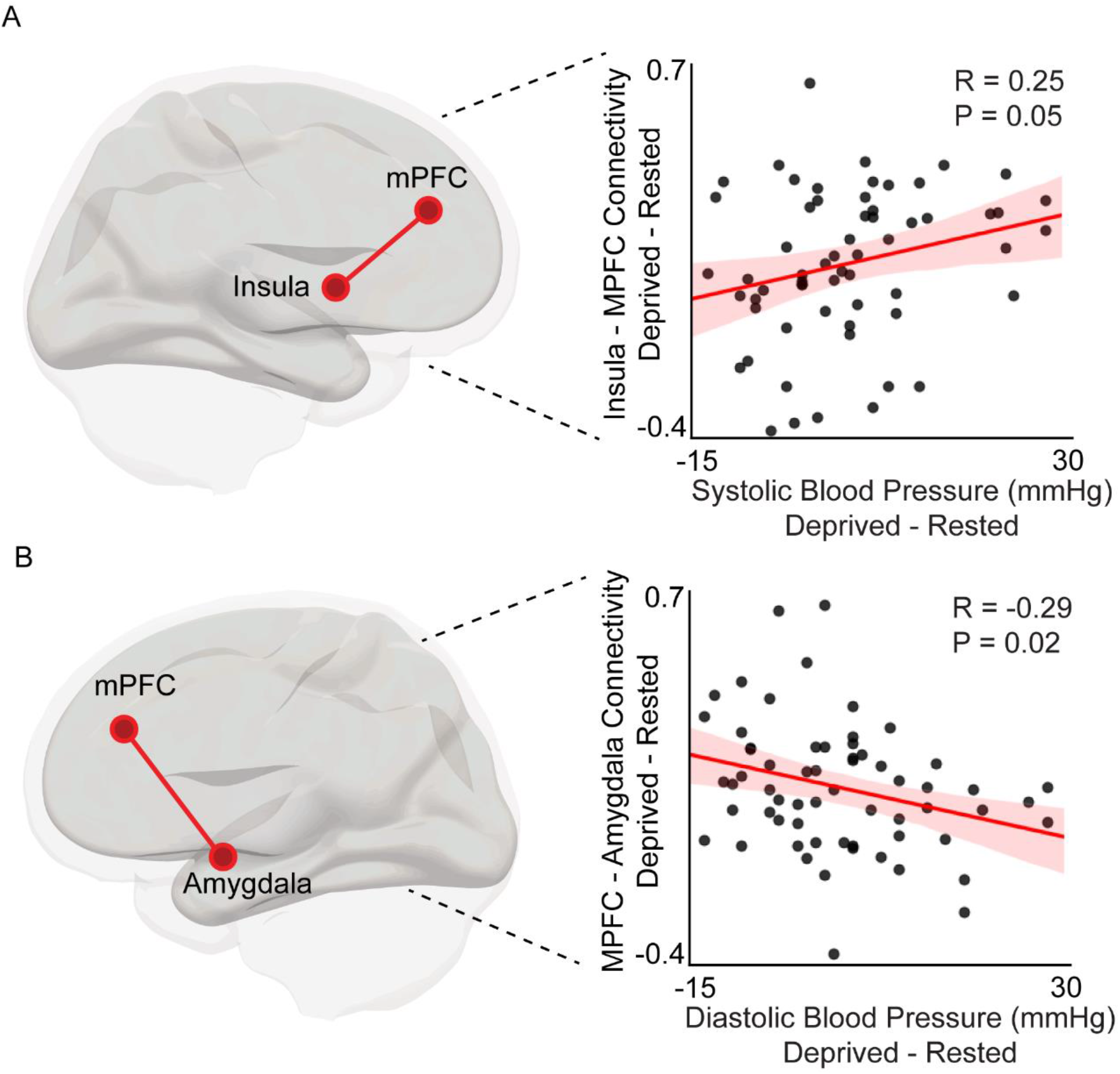
Brain connectivity and cardiovascular changes. **A**, Scatterplot represents significant positive correlation between sleep-loss related increases in insula and MPFC connectivity and sleep-loss related increases in systolic blood pressure. **B**, Scatterplot represents significant negative correlation between sleep-loss increases in MPFC and amygdala connectivity and sleep-loss related increases in diastolic blood pressure.

To better understand the specificity of central brain changes and cardiovascular changes, post-hoc analyses examined associations between sleep-loss related alterations in four non-a-priori, though well-recognized, larger-scale resting-state brain networks: the Default-Mode Network (DMN), Dorsal Attention Network (DAN), Ventral Salience Network (VSN), and Somatomotor Network (SMN)^33^. Neither differences in connectivity within or between these large-scale resting-state networks showed significant associations with the sleep-loss related increases in blood pressure, the decrease in heart rate, or the increase in heart rate variability (all p > 0.05). Therefore, local changes within the specific a priori cardiovascular control network^3,9-13^, rather than global changes in larger-scale brain networks^33^, best accounted for changes in peripheral blood pressure.

#### Emotion: Interactions with Brain Connectivity & Cardiovascular Function

Independent of sleep, mood and anxiety have well-recognized relationships with peripheral autonomic activity, including changes in cardiac and vascular activity^34^. Mood states and anxiety have also been related to changes in activity and functional connectivity of some nodes of the cardiovascular control network, specially the insular and MPFC nodes that are central to emotional brain processes^35-37^. Furthermore, both positive and negative mood states are deleteriously impacted by sleep loss^38^.

Building on these inter-connected links, we finally tested the hypothesis that changes in mood and anxiety caused by sleep loss would be significantly associated with, rather than independent of, the sleep-loss related changes in cardiovascular state and cardiovascular control brain networks.

As expected^38,39^, sleep deprivation was associated with a worsening of mood: a reduction in positive mood (t(65) = 6.8, p < 0.0001) and an increase in negative mood (t(65) = −5.2, p < 0.0001), relative to the sleep rested condition. Sleep loss also resulted in an increase in state anxiety, relative to the sleep rested state (t(65) = −4.3, p < 0.0001).

Demonstrating affective specificity, sleep-loss increases in negative mood state were associated with sleep-loss increases in blood pressure, such that the greater the negative mood increase, the greater the increase in diastolic blood pressure, though this was not significant following FDR correction (r(59) = 0.32, p = 0.18; FDR-corrected). Moreover, sleep-loss changes in heart rate and heart rate variability were not associated with changes in any of the affective measures, (all p > 0.24, FDR-corrected), once again demonstrating specificity to changes in vascular control, rather than heart contractility rate. The association between negative mood and systolic blood pressure was not significant (r(60) = 0.12, p = 0.51, FDR-corrected).

Sleep-loss related reduction in positive mood was instead significantly associated with the sleep-loss change in connectivity between the left MPFC and ventromedial frontal cortex (r(65) = 0.32, p = 0.01; and trend levels with changes in left insula cortex and right MPFC connectivity (r(65) = −0.24, p = 0.06). Connectivity changes in the MPFC and insula regions also demonstrated significant associations with sleep-loss related changes in systolic and diastolic blood pressure, fitting with the roles of these areas in both affective and cardiovascular regulation^3,11,14,35-37^. No associations were evident for changes in negative mood (all p > 0.26).

Counter to our experimental hypothesis, however, sleep-loss changes in anxiety were not significantly associated with changes in a priori brain vascular control networks (**Table S2**).

Therefore, a dissociable influence of mood was observed, with sleep-loss amplifications in negative mood state selectively associated with the amplifications in peripheral body blood pressure, and the loss of positive mood state was expressly associated with impairments in viscerosensory brain connectivity. Such findings lend further support to an embodied framework of sleep loss in which changes in brain, body, and mental state are mutually related in their association with the sleep-loss induced shift towards hypertension.

### Discussion

Taken together, the current study demonstrates that (1) experimental sleep loss significantly and selectively increases peripheral blood pressure, independent of any increase in heart rate, (2) sleep loss simultaneously compromises functional brain connectivity within regions that help regulate vascular tone—the viscerosensory network, (3) the sleep-loss related changes in viscerosensory functional brain connectivity and in peripheral vascular tone were significantly inter-dependent, with the changes in brain nodes explaining the vascular shift towards hypertension, and (4) sleep-loss-related reductions in positive state and amplification in negative state each showed a signification additive interactions with the selective impairments in brain network activity and peripheral hypertension caused by sleep loss, consistent with mental-state influence.

#### Sleep Loss and Cardiovascular State

Chronic sleep disturbances, including sleep disorders such as insomnia and sleep apnea, are recognized risk factors for hypertension and cardiovascular disease^1,2,4,5^. Moreover, experimental sleep loss causally increase blood pressure^6,7^. However, the underlying mechanism(s) by which poor sleep increases blood pressure are poorly understood.

The heart is known to receive both sympathetic and parasympathetic input, yet most vessels (arteries and veins) only receive sympathetic innervation^40^. Based on the dissociation observed, one parsimonious mechanism is that increases in blood pressure caused by sleep deprivation are triggered by an increase in sympathetic tone, while the decrease in heart rate reflects a separable shift in cardiac-mediated autonomic balance towards the influence of parasympathetic input^41^. That is, an autonomic dissociation caused by sleep loss, wherein increased sympathetic output dominates in peripheral vessels, yet the opposite is true in the heart. This possibility is also supported by the sleep-loss related increase in heart rate variability—a measure reflecting vagal (parasympathetic) modulation of the heart^31^. Indeed, the high sleep pressure resulting from sleep deprivation increases parasympathetic cardiac drive slowing heart rate^26,28^.

#### Sleep Loss & Resting Brain Connectivity

Peripheral functions of the body, including the regulation of the autonomic nervous system, are causally influenced by a set of brain regions involved in viscerosensory and cardio-modulatory processes^3,14,42^. The second component of our experimental hypothesis sought to determine whether changes in connectivity within specific brain networks that regulate cardiovascular state occurred under conditions of sleep deprivation, and whether such changes were related to the observed changes in cardiovascular function.

At a main effects level, sleep loss resulted in a collection of connectivity changes within our a priori set of brain regions, the majority of which showed increases in their nodal connectivity. These connectivity changes all jointly shared at least one of two nodes: the insula and MPFC. The insular cortex is not only sensitive to sleep status^43,44^, but is of central importance in the registration and control of bodily states, including cardiovascular state^3,11-13^. The MPFC, as with the insula, is also sensitive to sleep quality and quantity^39,45-47^, and orchestrates cardiovascular-challenge responses^13^.

Our findings corroborate and extend previous reports, indicating that sleep-loss-related changes in insula and MPFC connectivity may reflect impairment of brain networks involved in the coordination of cardiovascular physiology. Fitting this proposition, these brain regions are highly interconnected with several cortical, subcortical, and brainstem regions that dictate autonomic activity^3,11-14^.

In contrast to the dominant increases in connectivity caused by sleep loss, one pathway showed a decrease in connectivity under conditions of sleep loss: the link between the MPFC and ventromedial frontal cortex. Both regions have been previously associated with sympathetic nervous system drive and the central autonomic network of the brain^14^. Lesioning of the MPFC is associated with increased hypothalamic-pituitary-adrenal axis stress responses^17^, which is of central importance for the integration of brain and vascular function^18^. Furthermore, reduced activation in the MPFC has been associated with increased stress-induced vasoconstriction^15^.

#### Sleep Loss, Cardiovascular State & Resting Brain Connectivity

Adding to the findings of sleep-loss related changes in cardiovascular state and brain-network connectivity, we further demonstrate that these changes are significantly associated. First, the increase in systolic blood pressure was associated with increased connectivity between two regions involved in the regulation of vascular tone: the insula and MPFC. Both the insula and MPFC are involved in the representation and generation of visceral states, including cardiovascular reactions^48^. Thus, these two regions, along with a network of corticolimbic and brainstem systems, play instrumental roles in tuning cardiovascular and autonomic adjustments in response to contextual demands^13,42,48^. Indeed, the insula and MPFC are multi-synaptically connected to pre-autonomic nuclei which innervate both the heart and vasculature^49,50^. There is also evidence that increased resting activity in the insula and MPFC predicts greater evoked blood pressure reactions, interpreted as signaling increased sympathetic-mediated blood pressure reactivity^48^.

A different part of the cardiovascular control network was observed to be associated with the sleep-loss increase in diastolic blood pressure. Here, the magnitude of sleep-loss increase in diastolic blood pressure was significantly associated with decreased connectivity between the MPFC and amygdala. The amygdala is instrumental in coordinating behavioral and physiological adjustments through connections to cortical, hypothalamic, and brainstem nuclei involved in stress processing and autonomic-cardiovascular control^48^. Additionally, the amygdala can impact blood pressure through influence over the baroreflex, which serves to maintain blood pressure through autonomic regulation of heart rate, cardiac output, and vascular resistance, and is associated with preclinical atherosclerosis^9,51^. Thus, the reduction in amygdala connectivity to other regions involved in cardiovascular control (MPFC) could represent a degree of disorganization of brain homeostatic functions.

The vascular system is subject behavioral, autonomic, and endocrine regulation^52^. The complexity of these interacting pathways suggests that altered blood pressure control is due to a failure of a central control system, namely the brain^53^. Notably, in the current data, the decrease in heart rate was not associated with the increase in blood pressure. Furthermore, blood pressure-related changes in functional connectivity in the brain were similarly not associated with heart rate. Taken together, it is likely that multiple, perhaps counteractive, processes underlie the conflicting vascular and cardiac responses to sleep loss, fitting with a loss of central brain regulation.

#### Sleep Loss, Mood States, Brain, and Body

The final component of the study examined inter-relationships with changes in mood state and anxiety. Independent lines of research have demonstrated that sleep-loss detrimentally impacts affective states, including mood^38^. Moreover, mood states are known to causally alter cardiovascular function, including peripheral blood pressure^34^. In addition, the regulation of positive and negative mood states is further controlled by overlapping brain regions to those that are known to form the basis of central brain cardiovascular control, notably the insula and MPFC^35-37^.

Fitting with prior findings^38,39^, sleep loss significantly worsened mood state through a reduction in positive mood and an increase in negative mood. We extend these findings, establishing that sleep-loss related increases in negative mood are further related to physiological changes within the body, here the shift towards hypertension.

Indicating potential mood-type specificity, we further show that, this affective link to visceral function was specific to peripheral-body vascular tone and was significant for the increase in negative mood, and not the decrease in positive mood. Instead, the link to visceral function and the sleep-loss reduction in positive mood was specific to the changes in functional brain connectivity, notably the viscerosensory network regions of the insula and MPFC.

These latter finding may therefore suggest a dissociable influence of affective mental state on vascular regulation under conditions of sleep loss: negative mood state linked to peripheral body vascular function, yet the loss of positive mood state linked to central brain visceral regulation. That there is significant overlapping functionality of the MPFC and insula in both emotional and viscerosensory brain regulation^35-37^ further suggests a convergence of regulatory function in these brain regions underpinning an embodied framework of sleep loss, one in which sleep deprivation impacts affective and cardiovascular function through impairment of core regulatory centers in the brain.

More generally, our results indicate the presence of a coupled relationship between vascular responses to sleep loss, brain function and mental-state changes in affect. Such findings impress the previously proposed conception of sleep loss triggering broad homeostatic biological distress caused by allostatic overlap^54^. Pragmatically, these findings highlight the target of sleep as a risk factor, and modifiable intervention target, determining cardiovascular disease risk, here through regulation of central brain and mental-state mechanisms that control cardiovascular function with the goal of maintaining vascular tone within a normotensive range.

### Methods and Materials

#### Experimental Model and Subject Details

Sixty-six healthy adults aged 18-24 years (mean ± SD: 20.7 ± 1.7, 52% female) completed a repeated-measures cross-over design (described below). There was no influence of sex on cardiovascular outcomes (p > 0.64). Participants abstained from caffeine and alcohol for 72 hours before and during the entire course of the study and kept a normal sleep-wake rhythm (7-9 hours of sleep per night) with sleep onset before 1:00 A.M. and rise no later than 9:00 A.M. for the three nights before the study participation, as verified by sleep logs and actigraphy.

Exclusion criteria, assessed using a prescreening semi-structured interview, were as follows: a history of previously-diagnosed sleep disorders, neurological disorders, closed head injury, Axis 1 psychiatric disorders according to the DSM-IV-TR criteria (encompassing mental disorders, including depression, anxiety disorders, bipolar disorder, attention deficit disorder, and schizophrenia), history of drug abuse and current use of antidepressant or hypnotic medication, nicotine use, consumption of more than five alcoholic drinks per week, crossing of time zones in the 3 months before the study, and general contraindications to MRI. Participants also had blood pressures within normotensive ranges, confirmed with a blood pressure reading of less than 120/80 mmHg at the time of consenting^55^. Participants who reported sleeping <7 hours per night or consuming three or more daily caffeine-containing drinks were also excluded from entering the study. The Pittsburg Sleep Quality Index^56^ was employed to determine the quality of recent sleep history, and participants considered poor sleepers (global score > 5) were excluded. The study was approved by the UC Berkeley Committee for the Protection of Human Subjects, with all participants providing written consent.

#### Experimental Design

To test the experimental hypotheses, participants entered a repeated-measures study design (**Fig 1**), including two sessions—one following a normal night of sleep and one after 24 hours of total sleep deprivation. The two sessions were separated by at least 7 days (mean ± SD: 9.8 ± 3.8), with the order of the sleep-rested and deprived conditions counterbalanced across subjects.

In the sleep deprived session, participants arrived at the laboratory at 10:00 P.M. and were continuously monitored throughout the enforced waking period. Activities during the sleep deprivation condition were limited to use of the internet, email, short walks, reading, movies of low emotionality, and playing board games, thereby providing a standardized regimen of waking activity without undue stress. Participants also avoided exercise during the experimental sessions. In the sleep-rested session, participants came to the laboratory at 8:00 P.M. and were prepared for an 8-hour night of sleep (∼11:00 P.M. to 7:30 A.M. ± 30 min). In both conditions, mood and anxiety were assessed in the morning at the time of the cardiovascular assessment using the Positive and Negative Affective Scale and State-Trait Anxiety Inventory questionnaires, respectively^57,58^. The following morning, participants’ cardiovascular state was measured followed by fMRI scanning.

#### Cardiovascular Assessments

In each condition, cardiovascular state was assessed to determine blood pressure, heart rate, and heart rate variability. These assessments were performed approximately 1 hour after awakening, and at circadian-matched times in the sleep rested and sleep deprivation conditions for each participant.

Blood pressure measurement was performed with Omron BP742 blood pressure monitor (Omron Healthcare Europe BV, Hoofddorp, The Netherlands), fitted to the participants’ nondominant arm over the brachial artery. Participants were instructed to remain motionless with open eyes while sitting with feet flat on the floor during the recording with rested cuffed arm at heart height, consistent with standard blood pressure recording guidelines^59^. Two consecutive measurements were taken and the mean of these two measurements was used for analyses.

Immediately following blood pressure assessment, participants were fitted with electrodes for bipolar electrocardiography (ECG), and cardiac activity was recorded for five minutes. Electrodes were applied in a lead II format with the reference electrode placed below the right clavicle parallel to the right shoulder and a second electrode placed on the torso at the fourth intercostal space on the left side parallel to the left hip. Similar to the blood pressure recording, participants sat motionless with open eyes while sitting with feet flat on the floor. All data were screened for measurement artifacts and beat-to-beat intervals extracted using ARTiiFACT software^60^, and all flagged interbeat intervals were visually checked. If confirmed as artifactual, they were deleted and substituted by means of cubic spline interpolation of neighboring intervals. The time-domain measure pNN50 was calculated for each participant, which is the percentage of differences between adjacent interbeat intervals (NN) that are greater than 50ms and represent a recommended measure of vagally-mediated heart rate variability^31^.

#### fMRI preprocessing

Preprocessing was performed using fMRIPrep 1.2.5 (RRID:SCR_016216). The T1-weighted image was corrected for intensity non-uniformity using N4BiasFieldCorrection and used as a reference throughout the analysis. Spatial normalization to the ICBM 152 Nonlinear Asymmetrical template was performed through nonlinear registration with antsRegistration using brain-extracted version of both T1-weighted volume and template (ANTs 2.2.0 RRID:SCR_004757). Brain tissue segmentation of cerebrospinal fluid (CSF), white-matter (WM) and gray-matter (GM) was performed on the brain-extracted T1-weighted image using fast (FSL 5.0.10, RRID:SCR_002823).

For fMRI data, the mean BOLD image was co-registered to the T1-weighted reference using flirt. Head-motion parameters with respect to the BOLD reference (transformation matrices, and six corresponding rotation and translation parameters) were estimated before any spatiotemporal filtering using mcflirt. BOLD runs were slice-time corrected using 3dTshift from AFNI (RRID: SCR_005927), and the BOLD time-series (including slice-timing correction when applied) were resampled to MNI152NLin2009cAsym standard space. Known time-series factors requiring covariate accommodation were calculated based on the preprocessed BOLD: framewise displacement (FD), DVARS and three region-wise global signals. The three global signals are extracted within the CSF, the WM, and the whole-brain masks. Additionally, a set of physiological regressors were extracted to allow for component-based noise correction (CompCor^61^). Finally, regression was used to estimate and remove the whole-brain global signal time series, and functional images were smoothed using a 6 mm FWHM Gaussian kernel.

#### Statistical and fMRI analysis

Connectivity analyses were performed using the CONN toolbox version 19b (www.nitrc.org/projects/conn, RRID:SCR_009550). First, denoising was performed by regressing the six motion parameters, FD, and physiological noise components. The resulting BOLD time series was band-pass filtered (0.008 – 0.09 Hz), followed by region of interest (ROI) analysis, specifically ROI-to-ROI connectivity analysis.

Based on the experimental hypotheses, analyses focused on an a priori set of brain ROIs known to be involved central brain cardiovascular control^3,9-14^. These ROIs were drawn from anatomical atlases (Harvard-Oxford Cortical and Subcortical Atlas) included in the CONN toolbox, and were extracted using an ICA approach. For each ROI, the connectivity to all other ROIs was calculated for each subject in each condition separately, generating a symmetric correlation matrix wherein each element is the correlation between two ROIs in each condition and each participant.

In addition to ROI-to-ROI connectivity analyses, control analyses for specificity examined connectivity within and between additional prototypical brain networks of the Default-Mode Network (DMN), Dorsal Attention Network (DAN), Ventral Salience Network (VSN), and Somatomotor Network (SMN)^33^. Within-network connectivity was calculated for each network by extracting the mean resting-state BOLD time-series from each voxel within a region of a network and calculating the mean Pearson correlation with all other regions of the network. Between-network connectivity was similarly calculated for each network by extracting the mean resting-state BOLD time-series from every voxel within a network, and then calculating the correlation with the mean time-series of all other networks.

The resulting correlation coefficients from these fMRI analyses across participants in each condition were then statistically compared (two-tailed paired t-test) to determine the effect of sleep loss on brain connectivity. Consistent with imaging recommendations to correct for multiple statistical comparisons among the eight ROIs and four networks, the Benjamini-Hochberg procedure for controlling FDR was applied, with q = 0.05^32^. This procedure addresses the problem of simultaneously performing multiple significance tests by using an adaptive stepwise procedure to control the expected ratio of erroneous rejections to the number of total rejections. Only connectivity changes that survived the FDR criterion after correction were considered statistically significant and used in analyses. Pearson correlations were employed to test the relationships between sleep loss-related changes in brain connectivity and sleep loss-related changes in cardiovascular state. All statistical analyses were performed in Python using the Pingouin package^62^.

## Acknowledgments

We thank Olivia G. Murillo for her assistance in running the study.

## Author Contributions

AJK, RV, EBS, and MPW designed research. AJK, RV, and EBS performed research. AJK analyzed data. AJK and MPW wrote the paper with contributions from all authors.

## Declaration of Interests

Dr. Walker serves as a consultant for and has equity interest in Bryte, Oura, Shuni, and StimScience.

## Supplementary Materials

**Supplementary Table S1.**
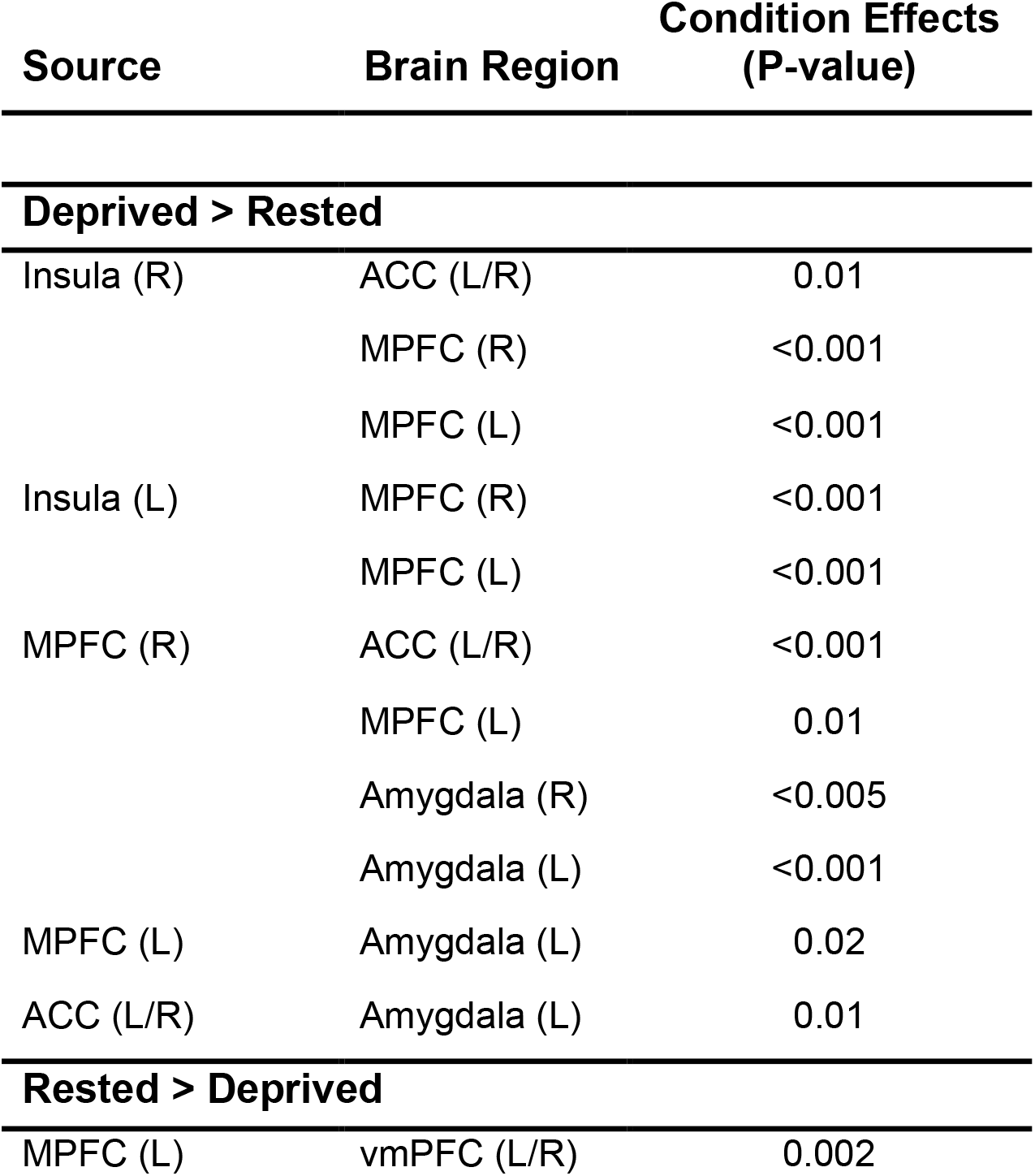
Resting Brain Connectivity. Significant condition effects (sleep rested < > sleep deprived) on brain connectivity, with all condition effects shown significant following FDR-correction. ACC, anterior cingulate cortex; MPFC, medial prefrontal cortex; vmPFC, ventromedial prefrontal cortex; HR, heart rate; HRV, heart rate variability (pNN50).

**Supplementary Table S2.**
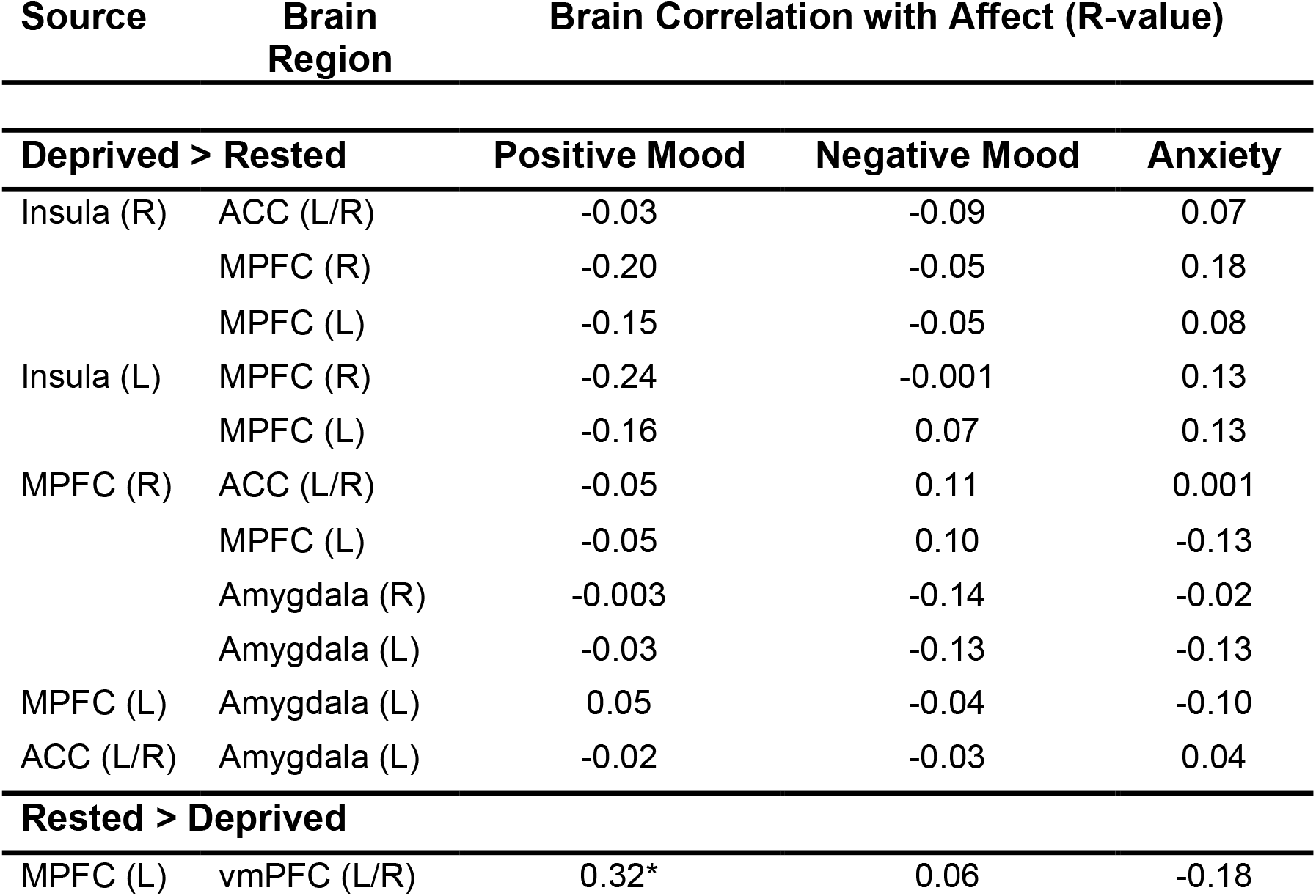
Connectivity Associations with Affect. Columns show the Pearson correlation between sleep-loss related changes in brain connectivity and changes in affective state, including mood (positive and negative) and anxiety. * significant following FDR-correction. ACC, anterior cingulate cortex; MPFC, medial prefrontal cortex; vmPFC, ventromedial prefrontal cortex.

| Source | Brain Region | Brain Correlation with Affect (R-value) |  |  |
| --- | --- | --- | --- | --- |
| Deprived > Rested |  | Positive Mood | Negative Mood | Anxiety |
| Insula (R) | ACC (L/R) | -0.03 | -0.09 | 0.07 |
|  | MPFC (R) | -0.20 | -0.05 | 0.18 |
|  | MPFC (L) | -0.15 | -0.05 | 0.08 |
| Insula (L) | MPFC (R) | -0.24 | -0.001 | 0.13 |
|  | MPFC (L) | -0.16 | 0.07 | 0.13 |
| MPFC (R) | ACC (L/R) | -0.05 | 0.11 | 0.001 |
|  | MPFC (L) | -0.05 | 0.10 | -0.13 |
|  | Amygdala (R) | -0.003 | -0.14 | -0.02 |
|  | Amygdala (L) | -0.03 | -0.13 | -0.13 |
| MPFC (L) | Amygdala (L) | 0.05 | -0.04 | -0.10 |
| ACC (L/R) | Amygdala (L) | -0.02 | -0.03 | 0.04 |
| Rested > Deprived |  |  |  |  |
| MPFC (L) | vmPFC (L/R) | 0.32* | 0.06 | -0.18 |

## Notes

### Competing Interest Statement

The authors have declared no competing interest.

